# Protein Contaminants Matter: Building Universal Protein Contaminant Libraries for DDA and DIA Proteomics

**DOI:** 10.1101/2022.04.27.489766

**Authors:** Ashley M. Frankenfield, Jiawei Ni, Mustafa Ahmed, Ling Hao

## Abstract

Mass spectrometry-based proteomics is constantly challenged by the presence of contaminant background signals. In particular, protein contaminants from reagents and sample handling are often abundant and almost impossible to avoid. For data-dependent acquisition (DDA) proteomics, exclusion list can be used to reduce the influence of protein contaminants. However, protein contamination has not been evaluated and is rarely addressed in data-independent acquisition (DIA). How protein contaminants influence proteomics data is also unclear. In this study, we established protein contaminant FASTA and spectral libraries that are applicable to all proteomic workflows and evaluated the impact of protein contaminants on both DDA and DIA proteomics. We demonstrated that including our contaminant libraries can reduce false discoveries and increase protein identifications, without influencing the quantification accuracy in various proteomic software platforms. With the pressing need to standardize proteomic workflow in the research community, we highly recommend including our contaminant FASTA and spectral libraries in all bottom-up proteomics workflow. Our contaminant libraries and a step-by-step tutorial to incorporate these libraries in different DDA and DIA data analysis platforms can be valuable resources for proteomics researchers, which are freely accessible at https://github.com/HaoGroup-ProtContLib.

## INTRODUCTION

Mass spectrometry (MS)-based proteomics is constantly challenged by exogenous contaminants and interferences that can be introduced into samples throughout the experimental workflow. Contaminations from polymers, detergents, solvents, ion sources, and other additives are often singly charged, which can be avoided by the careful selection of reagents or removed by ion mobility MS interface *(e.g.,* FAIMS).^1–3^ However, contaminant proteins and peptides are almost impossible to eliminate from the experimental workflow. For example, keratins from researchers’ skin and hair can be found on all surfaces and dust during sample handling.^4^ Rodent and sheep keratins can originate from animal facilities and wool clothing. Residue cell culture medium can lead to bovine protein contaminations. Protein digestion enzymes (*e.g*., trypsin and Lys-C) and the production of enzymes can introduce protein contaminants into bottom-up proteomics workflow.^1^ Additionally, bovine serum albumin (BSA), immobilized antibodies, and affinity tags *(e.g.,* streptavidin, FLAG, HA) from affinity columns/beads also represent major contaminants in immunoassays and affinity purification MS.^5,6^ These exogenous contaminant proteins/peptides can compete with real samples in the MS ion source, occupy the cycle times in the mass analyzer, reduce the number of useful peptide spectra, and hinder the detection of low abundant proteins from complex biological samples.

Sample type-specific interferences have been evaluated previously and are important proteomic resources, such as non-specific interactions in affinity purification and contaminations from plasma proteomics.^6,7^ It is important to note that these interference proteins are specific to certain types of experiments and may actually be useful proteins in other types of proteomic studies. Therefore, these interference proteins cannot be marked as universal exogenous contaminant proteins for all proteomic experiments. Due to the negative effects of protein contaminants in MS-proteomics, various methods have been implemented to combat this problem. Keratin contaminations can be reduced by using a laminar flow hood and fastidiously wiping down surfaces with ethanol and water.^8^ However, it is almost impossible to eliminate keratins from proteomic experiments. Contaminations from proteolytic enzymes and affinity tags can be reduced by carefully optimizing the amount of enzymes and beads. Nevertheless, such practices may not be feasible for MS facilities and biological samples with limited amounts. For data-dependent acquisition (DDA) proteomics, an exclusion list can be used to disregard specific ions from being isolated for MS/MS fragmentation.^9–12^ However, an exclusion list is highly specific to LC-MS gradient and instrumentation, which is difficult to transfer across different MS platforms and laboratories. Peptides with similar *m/z* and retention times could also be accidentally excluded in complex biological samples. Contaminant peptides are not entirely useless and can be used as quality control to evaluate sample preparation reproducibility. Trypsin peptides can also be used to normalize retention time.^13^ In order to mark contaminant proteins from the dataset, contaminant FASTA libraries can be used in various DDA software platforms.^14–18^ The most widely used contaminant FASTA files are from MaxQuant^15^ and cRAP (https://www.thegpm.org/crap/). However, these FASTA files have not been updated in years and contain many deleted/unassigned UniProt entries and human protein standards which are not contaminant proteins.

Despite various strategies to reduce the influence of protein contaminants in DDA proteomics, protein contamination has not been evaluated and is rarely addressed in data-independent acquisition (DIA) proteomics. Many exogenous contaminants from different species cannot be identified unless included in FASTA or spectral libraries. DDA exclusion list is not compatible with DIA because all co-eluting peptides within a pre-determined isolation window are fragmented together regardless of precursor intensities in DIA. Due to the wide isolation window in DIA, we hypothesized that contaminant proteins/peptides could be especially problematic if left unaddressed, leading to false identifications in DIA proteomics.

DIA data analysis can be conducted using spectral library-based software tools *(e.g.,* OpenSWATH^19^, Spectronaut^20^, DIA-NN^21^, Skyline^22^, EncylopeDIA^23^, MaxDIA^24^) or library-free strategies with in-silico digested pseudo peptide spectra based on FASTA protein sequences *(e.g.,* DirectDIA^25^, DIA-NN^21^, DIA-Umpire^26^, PECAN^27,28^, DeepDIA^29^). While contaminant FASTA libraries are widely implemented in DDA data analysis, they are rarely used for DIA data analysis.^13,30^ The PRIDE^31^ website provided a contaminant spectral library based on the commonly used cRAP list. However, the cRAP contaminant list has not been updated for 10 years and contains many noncontaminant human protein standards such as cathepsins, annexin, and myoglobin.

In this study, we created a series of contamination-only samples to establish the universal contaminant protein spectral and FASTA libraries that can be used in all bottom-up proteomic experiments. We then evaluated how protein contaminants and contaminant libraries influence identification and quantification in DDA and DIA proteomics. The benefits and applicability of these contaminant libraries were demonstrated in various DDA and DIA data analysis platforms. These contaminant FASTA and spectral libraries are freely accessible at https://github.com/HaoGroup-ProtContLib with a step-by-step user manual to promote standardized and reproducible proteomics data analysis and reporting pipeline in the broad proteomics community.^32–34^

## MATERIALS AND METHODS

### Generation of Contaminant-Only Samples

We generated a series of contaminant-only samples by adding different proteolytic enzymes to the lysis buffer (1M Urea in 50 mM Tris-HCl), commonly used beads coated with affinity tags, and fetal bovine serum (FBS) that is commonly used for cell culture medium.The proteolytic enzymes used here include sequencing grade Trypsin (V5111), Trypsin Gold (V5280), Trypsin/Lys-C (PRV5073), and Lys-C (VA1170) from Promega. The beads used here include Sero-Mag streptavidin magnetic beads (Cytivia), Anti-Flag M2 affinity agarose beads (Sigma), and EZview Red anti-HA affinity agarose beads (Sigma). Clean ungloved hands were purposely rubbed together above these samples to increase keratin contaminations.

### Human Cell Culture and Mouse Brain Tissues

HEK293 cells were maintained in DMEM/F12 HEPES medium containing 10% of FBS. Mouse brain samples were obtained from wild-type mice (C57/B6) under protocols approved by the George Washington University Institutional Animal Care and Use Committee. HEK cells and mouse brain samples were lysed in 8 M Urea in 50 mM Tris-HCl buffer and sonicated for 15 min in an ice-cold water bath using a QSonica Q700 Sonicator with alternating cycles of 1 min on and 30 s off. Protein lysates were clarified by 15 min of centrifugation at 12,000 rpm at 4 °C and stored in −80 °C. Total protein concentrations were determined using a detergent-compatible colorimetric protein assay (DCA, BioRad).

### Proteomic Sample Preparation

The routine bottom-up proteomic workflow was conducted for contaminant-only samples, HEK cells and mouse brain lysates as described previously.^35,36^ Briefly, disulfide bonds were reduced using 5 mM Tris(2-carboxylethyl)phosphine (TCEP) for 30 min, 15 mM of iodoacetamide for 30 min in dark, and 5 mM TCEP for 10 min on a ThermoMixer shaking at 1,200 rpm at 37 °C. Protein digestions were conducted using various enzymes (contaminant-only samples) and Trypsin/Lys-C (HEK and mouse samples) for 18 hours at 37 °C on a ThermoMixer, and quenched with 10% trifluoroacetic acid until pH < 3. Peptides were then desalted on a Waters Oasis HLB Plate using the manufacturer’s protocol, dried down under SpeedVac, and stored at −30 °C.

### LC-MS/MS Analysis for DDA and DIA Proteomics

Peptide samples were analyzed on a Dionex UltiMate 3000 RSLCnano system coupled with a Thermo Fisher Q-Exactive HF-X mass spectrometer. The mobile phase buffer A was 0.1% formic acid in water, and buffer B was 0.1% formic acid in acetonitrile. HEK cells and mouse brain samples were injected onto an Acclaim PepMAP C18 trap column (3 μm, 100Å, 75 μm × 2cm) and further separated on an Easy-spray PepMap C18 column (2 μm, 100Å, 75 μm × 75cm) with a flow rate of 0.2 μL/min, an LC gradient of 210 min, and a column temperature of 55 °C. Contaminant-only samples were analyzed with a 15 cm PepMap C18 column, a flow rate of 0.3 μL/min, and an LC gradient of 120 min. For DDA analysis, MS scans from *m/z* 380 to 1,500 with a resolving power of 120K (at *m/z* 200 FWHM), an automatic gain control (AGC) target of 1 × 10^6^, and a maximum injection time (maxIT) of 50 ms. Precursors were isolated at a window of *m/z* 1.4 and fragmented with a normalized collision energy (NCE) of 30%, a resolving power of 7.5K for MS/MS, and a maxIT of 40 ms. For DIA analysis, MS scans from *m/z* 400 to 1000 at a resolving power of 60K, an AGC target of 1 × 10^6^, and a maxIT of 30 ms. The precursor isolation window was set to *m/z* 8.0 (staggered) with 75 sequential DIA MS/MS scans between *m/z* 400 to 1000 at a resolving power of 30K, an AGC target of 5 × 10^5^, a MaxIT of 30 ms, and an NCE of 30%.

### Repository Data from ProteomeXchange

Two repository datasets from the ProteomeXchange website were downloaded and reanalyzed using our contaminant libraries. Repository dataset A is a HepG2 human cell DIA dataset (PXD022589) containing 27 raw data.^24^ Dataset B is a fractionated mouse cortex DIA dataset (PXD005573) containing 12 raw data.^37^ Additionally, a fractionated HEK and HeLa cell DDA dataset (PXD001468) was used to generate a spectral library for library-based DIA data analysis.^38^

### DDA Proteomics Data Analysis

All DDA proteomic datasets in this study were analyzed with both the MaxQuant (2.0.2.0) and Thermo Fisher Proteome Discoverer (2.4.1.15) software. Contaminant-only samples were analyzed with the new contaminant FASTA library only. HEK cells and mouse brain samples were analyzed using the Swiss-Prot *Homo sapiens* database (reviewed) and *Mus musculus* database (reviewed), respectively, with and without our contaminant FASTA library. The false discovery rate (FDR) cutoff for protein and peptide spectral matches (PSMs) identifications was set at 0.01. Trypsin or Lys-C enzyme was used with a maximum missed cleavage of two. Precursor tolerance was set to 20 ppm. The fixed modification was cysteine carbamidomethyl, and variable modifications were methionine oxidation and protein N-terminus acetylation.

### DIA Proteomics Data Analysis

#### Spectronaut software

Several spectral libraries were generated using Pulsar in Spectronaut 15 with “BGS Factory settings”.^20^ The contaminant spectral library was generated using the set of contaminant-only DDA dataset. Two spectral libraries were generated for each sample type (mouse brain and human cell) with and without including the contaminant-only samples (**Supplemental Table S2**). Specific trypsin digestion was set with a maximum of two missed cleavages. A fixed carbamidomethyl modification of cysteine, and up to three variable modifications for oxidation of methionine and acetylation of the protein N-terminus were allowed. PSM, peptide and protein FDR were set to 0.01. Both library-based and library-free (DirectDIA) analyses were performed in Spectronaut 15 using default settings. The quantification step was modified to perform an interference correction that used only identified peptides to train the machine-learning model. No cross-run normalization or imputation of missing values was used.

#### DIA-NN software

DIA-NN (v1.8) was used for both spectral library-based and library-free DIA analyses.^21^ Raw data files were converted to the open-format *.mzML* using the msConvert feature of the ProteoWizard package.^39^ Library-based analysis was conducted in DIA-NN using the spectral libraries established above in Spectronaut Pulsar. A fixed carbamidomethyl modification of cysteine, and up to three variable modifications for oxidation of methionine and acetylation of the protein N-terminus were allowed. Protein interferences were removed based on gene ID. FDR (0.01) was controlled by manually filtering the protein and peptide q-values in the report file. For library-free analysis, FASTA digest was selected. The spectral libraries were also included to train the deep learning model.

### Post-Data Analysis Filtering

To increase the confidence of protein/peptide identifications, proteins that were identified with only one precursor or an intensity below 10 were removed from all datasets using R. Contaminant proteins can be easily filtered out from the results by searching the “Cont_” prefix in the UniProt ID column from the result files. Contaminant proteins were removed before calculating the coefficient of variation and Spearman’s correlation to evaluate proteomics quantification.

### Data Availability

All raw files have been deposited to the ProteomeXchange Consortium with the data identifier, PXD031139. The protein contaminant library and step-by-step user tutorial are also freely accessible at https://github.com/HaoGroup-ProtContLib.

## RESULTS AND DISCUSSION

### Building the Contaminant Protein FASTA and Spectral Libraries

Most exogenous contaminant proteins originated from reagents and sample handling are commonly shared in all bottom-up proteomic experiments. Therefore, we aim to build universal contaminant protein libraries that can be used in all bottom-up proteomics (**Figure 1**). Although widely used for DDA proteomics, protein contaminant lists from MaxQuant and cRAP have not been updated for years, containing many deleted/unassigned UniProt IDs, sample-specific interference (noncontaminant) proteins, and commercially available human protein standards which are incorrectly listed as contaminant proteins. Therefore, we first built a new contaminant FASTA library by manually merging the available contaminant lists online, updating their UniProt entry IDs, deleting noncontaminant proteins, searching for new contaminant proteins on UniProt, and combining them into a new FASTA file. Our new contaminant FASTA library contains 381 contaminant proteins including all human keratins related proteins, bovine contaminants from cell culture medium and affinity columns, various proteolytic enzymes, affinity tags, and other contaminants (**Supplemental FASTA and Table S1**). When compared to the MaxQuant and cRAP contaminant lists, our new FASTA library contains an additional 166 contaminant proteins (**Figure 2A**). This new FASTA library can be used for both DDA and DIA proteomics. We also added a “Cont_” prefix in each contaminant entry in the FASTA library, allowing contaminant proteins to be easily filtered and removed in the result files.

**Figure 1:**
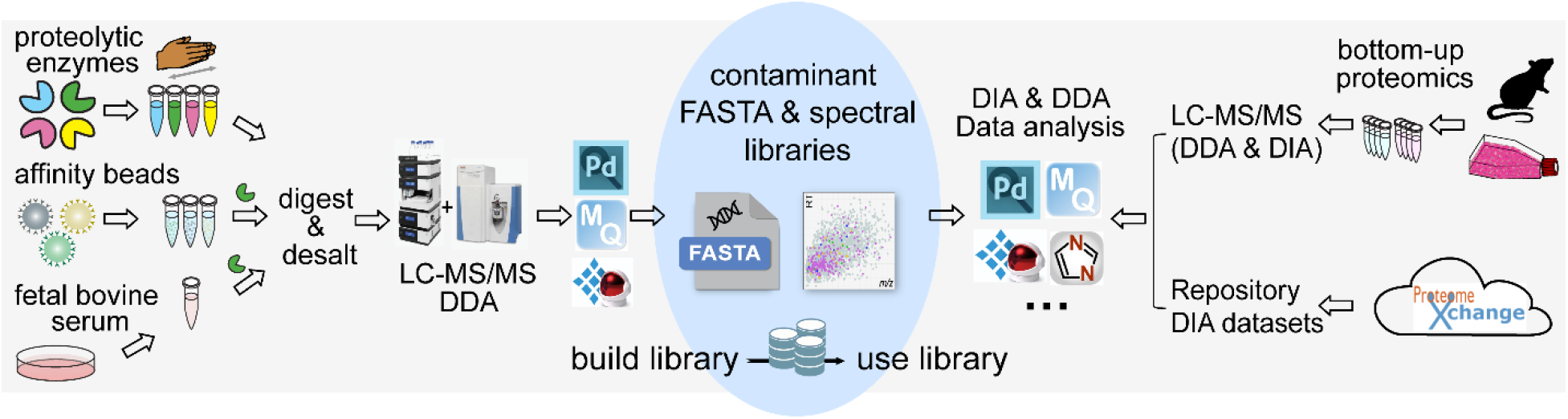
Schematic of building and using the contaminant libraries for DDA and DIA proteomics. A series of contaminant-only samples were created by adding different proteolytic enzymes to keratin-contaminated lysis buffer, commonly used beads coated with affinity tags, and fetal bovine serum (FBS) for cell culture medium. New contaminant FASTA and spectral libraries were created using DDA proteomic analyses of contaminant-only samples. These new contaminant libraries were evaluated using different biological samples and repository datasets in various DDA and DIA software platforms.

**Figure 2:**
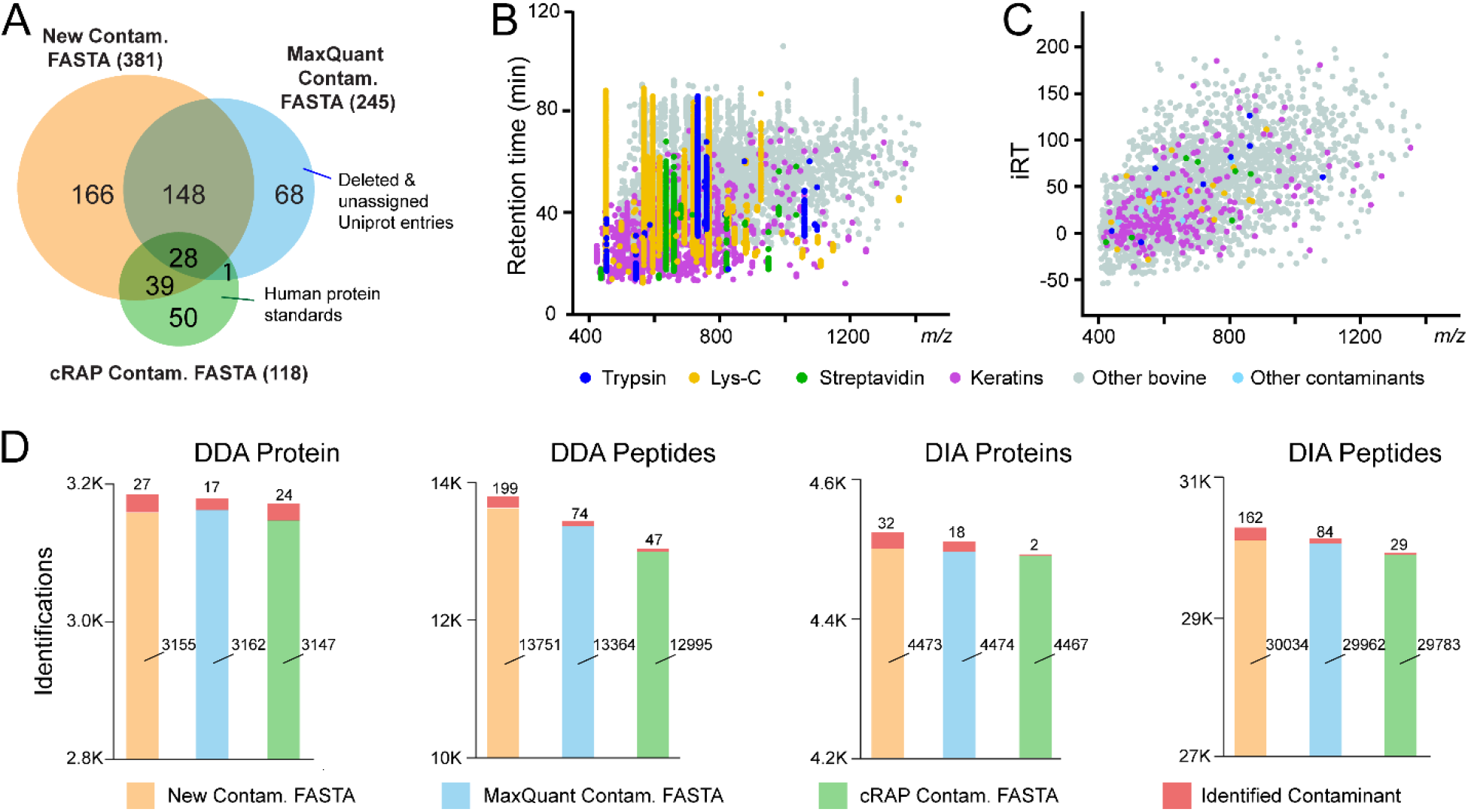
Characterization of the contaminant protein FASTA and spectral libraries. (A) Venn diagram comparison of contaminant protein lists from our newly generated contaminant FASTA and commonly used MaxQuant and cRAP contaminant FASTA files. (B) Scatterplot of identified contaminant peptides merged from contaminant-only samples in DDA LC-MS/MS analyses. (C) Scatterplot of contaminant peptides in our contaminant spectral libraries, generated by the Spectronaut Pulsar. iRT stands for in-silico normalized retention time. (D) Comparison of DDA and DIA protein and peptide identifications from HEK samples using our new contaminant FASTA in comparison to the MaxQuant and cRAP FASTA libraries.

To establish comprehensive contaminant protein spectral libraries for DIA proteomics, we created a series of contaminant-only samples using various proteolytic enzymes, affinity purification beads and fetal bovine serum (FBS) that are commonly used for cell culture medium. We validated the presence of each contaminant peptides by creating spectral libraries in MaxQuant, Proteome Discoverer and Spectronaut Pulsar. Hundreds of contaminant peptides were detected throughout the LC-MS gradient (**Figure 2B and Supplemental Table S3)**. Since trypsin and Lys-C are the two most commonly used enzymes for bottom-up proteomics, we created two DIA spectral libraries using Spectronaut Pulsar: tryptic contaminant peptides, and Lys-C-digested contaminant peptides. These spectral libraries are built from highly confident fragment ions assigned to each peptide sequence (**Figure 2C, Supplemental Table S4**), also freely accessible on ProteomeXchange (PXD031139). We compared our new FASTA library to the existing contaminant FASTA from MaxQuant and cRAP using DDA and DIA analyses of HEK samples. Improved protein/peptide identifications were achieved using the new library (**Figure 2D**). Further assessment of the contaminant proteins showed that fetal bovine serum proteins, human keratins and Lys-C enzyme produced the largest number of contaminant PSMs. Lys-C enzyme provides higher cleavage efficiency at lysine and is therefore often used in combination with trypsin to improve digestion efficiency.^40^ However, Lys-C enzyme contains almost two fold more arginine/lysine residues compared to trypsin, leading to many contaminant peptides. Additionally, bovine protein contaminants (albumin, *etc.)* were identified in all affinity purification beads despite conducting pre-washing steps. Streptavidin coated beads generated overwhelming streptavidin peptide signals.^5^ These exogenous contaminant proteins originated from a different species will not be identified unless the contaminant FASTA library is in use. Although our libraries provided commonly observed contaminant proteins for most proteomics experiments, contaminant proteins could be sample-specific. For example, keratins may be biomarkers for skin and oral cancer.^41^ In this special case, keratins may be important proteins that should not be removed. Our new FASTA library marked these common contaminant proteins with “Cont_” in the UniProt ID and we suggest researchers examine these proteins IDs before removing them from the results.

### Contaminant Peptides can Cause False Discoveries in DIA Proteomics

Contaminant FASTA library has been widely used for DDA proteomics, but is rarely included in DIA data analysis.^30,35,42,43^ Since DIA uses a much wider precursor isolation window (4-15 Da) compared to DDA (0.4-2 Da), contaminant peptides in DIA are more likely to be coeluted and cofragmented with other peptides. If not addressed properly, contaminant peptides may cause false identifications of peptides/proteins. To evaluate the influence of contaminant peptides, we analyzed several DIA proteomic datasets with and without our contaminant FASTA library. As shown in **Figure 3A**, when the contaminant FASTA library is not included during data analysis, a contaminant Lys-C peptide was misidentified as a KIF20B peptide due to numerous shared peptide fragments. After including the contaminant library, the peak picking algorithm identified an additional y3 ion and y7^++^ ion and assigned the fragmentation spectra to Lys-C instead of KIF20B with higher confidence and lower peptide *q*-values. This misidentification occurs frequently when Trypsin/Lys-C or Lys-C is used in multiple samples both during library generation and data processing (**Supplemental Figure S1A**). A similar scenario happened to a bovine contaminant protein SERPINA1 which was misidentified as CFAP100 (**Figure 3B**). Including the contaminant library allows the identification of three additional fragments to correctly assign to SERPINA1 contaminant peptide. We carefully examined the identification spectra in all datasets and found that these misidentifications do not happen on a large scale, yet still represent clear evidence of false discoveries caused by contaminant peptides when contaminant protein libraries are not in use. Furthermore, as contaminant peptides elute throughout the LC gradient and mass range (**Figure 2B**), many contaminant peptides can be coeluted and co-fragmented with real peptides of interest (**Supplemental Figure S1B-D).** Although co-elution and co-fragmentation are common in DIA proteomics, highly abundant contaminant peptides can suppress the detection of low abundant peptides by competing with them in the ion source and mass analyzer. In proteomics, a target-decoy strategy is commonly used to estimate the false discovery rate. High abundant contaminant peptides can generate high scores, potentially hindering the selection of low-score biologically meaningful proteins.^44^ Therefore, carefully optimizing experimental workflow to reduce contaminant signals and integrating contaminant libraries into the data analysis pipeline should be combined together to improve proteomics data quality.

**Figure 3:**
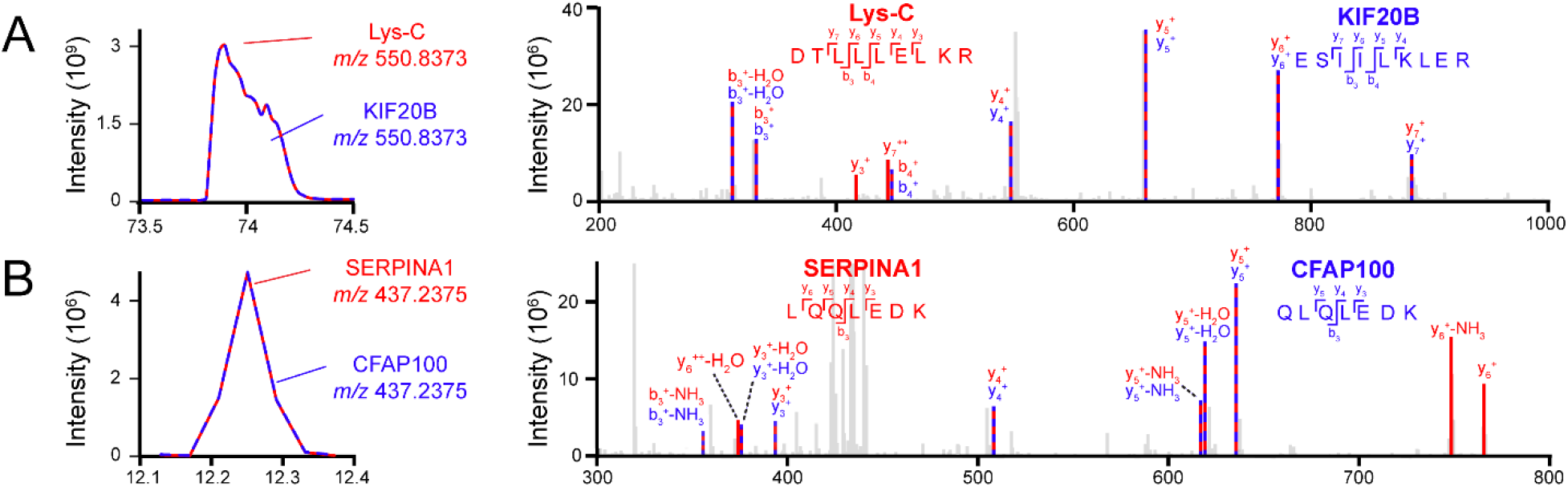
Examples of protein false identifications caused by contaminant peptides when contaminant library is not used in DIA data analysis. Example contaminant peptide chromatograms and MS/MS fragments were shown in red, real peptides of interest were shown in blue. (A) Lys-C (proteolytic enzyme contaminant) was misidentified as KIF20B. (B) SERPINA1 (bovine contaminant) was misidentified as CFAP100.

### Including Contaminant Protein Library Improves both DDA and DIA Proteomics

Contaminant libraries can be integrated into the DDA and DIA data analysis workflow via different strategies. DDA and library-free DIA analyses only require the contaminant FASTA protein sequences. Library-based DIA analysis requires both FASTA and spectral libraries. Contaminant spectral library can be generated in two ways: 1) an integrated spectral library built from contaminant-only raw data and custom proteomics data together; 2) two separate spectral libraries for contaminant and custom proteomics data. Contaminant FASTA file is also required when building these spectral libraries. In Spectronaut software, multiple spectral libraries can be included during data analysis. We found that the integrated spectral library performs similarly to two separate libraries with slightly higher total protein identifications in some datasets (**Supplemental Figure S2**). Either method is better compared to the results analyzed without the contaminant library. However, many other DIA software platforms do not allow the inclusion of multiple spectral libraries, and thus require an integrated spectral library. Including the additional contaminant FASTA and spectral libraries did not increase the software processing time for multiple DDA (Proteome Discoverer, MaxQuant) and DIA (DIA-NN, Spectronaut) platforms.

To demonstrate the benefits of contaminant protein libraries for both DDA and DIA proteomics, HEK cells and mouse brain samples were analyzed in DDA and DIA workflows in various data analysis software (**Figure 4**). After removing the contaminants, more peptides/proteins were identified when contaminant libraries were in use. The overall increase of protein and peptide identifications were around 0.9% and 1.3%, respectively, across all software and sample types. The improvement in noncontaminant protein IDs is likely due to decreased false identifications and altered target/decoy ratios when including the contaminant libraries. For DDA data, including contaminant FASTA improved noncontaminant peptide identifications but protein IDs were not influenced. Benefited from the additional contaminant spectral library, library-based DIA platforms provided a greater increase of identifications compared to library-free platforms. This is likely due to the high quality and abundant contaminant peptide spectra from our contaminant-only samples. For various DIA platforms, library-free DIA-NN generated the highest number of protein and peptide IDs possibly due to the deep learning model implemented in search algorithm and interference correction algorithm. Besides in-house generated proteomics data, we also analyzed repository datasets with and without contaminant library. Two DIA repository datasets were downloaded from ProteomeXchange: repository dataset A from HepG2 human cell samples^24^ and dataset B from mouse brain samples^37^. An increased number of proteins and peptides were identified when the data was analyzed with contaminant libraries (**Figure 5**). Particularly for repository dataset A, more than 5% of additional proteins and peptides (noncontaminants) were identified when contaminant library was used in library-based Spectronaut platform. Many bovine contaminant proteins were identified from repository dataset A, similar to our in-house generated HEK cell dataset, which can be traced back to the FBS used for human cell culture. To minimize the contaminations from cell culture medium, we highly recommend 2-3 times quick washes with phosphate-buffered saline (PBS) during cell harvest.

**Figure 4:**
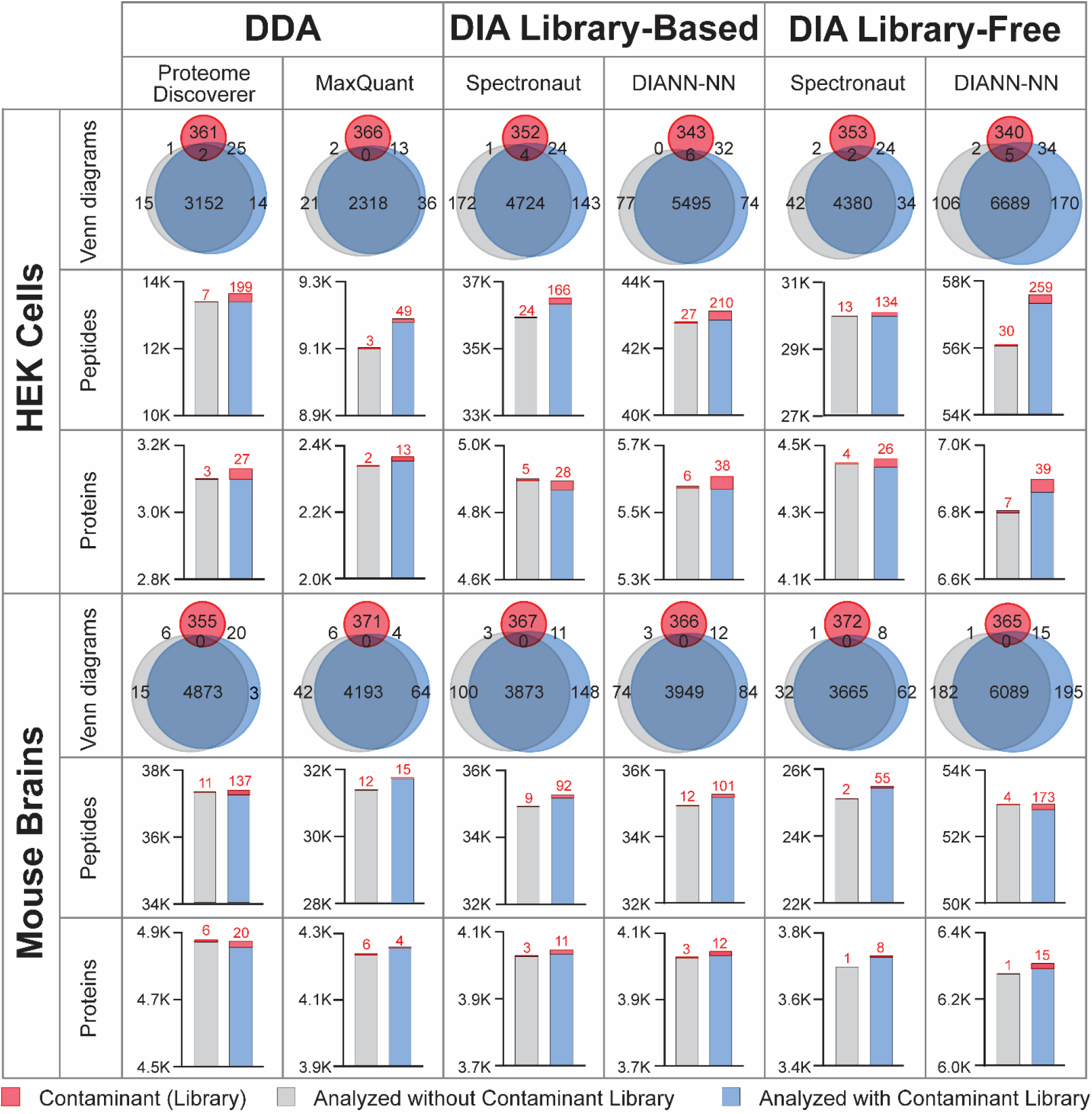
Evaluation of protein/peptide identifications influenced by the protein contaminant libraries in DDA and DIA proteomics. HEK cells and mouse brain samples were analyzed by various DDA and DIA software platforms, with (blue) and without contaminant libraries (grey). Venn diagrams showed the identified proteins from various datasets overlapping with the contaminant lists in the FASTA library (red). Bar graphs showed the identified contaminants (red) and noncontaminant proteins/peptides with and without using contaminant libraries.

**Figure 5:**
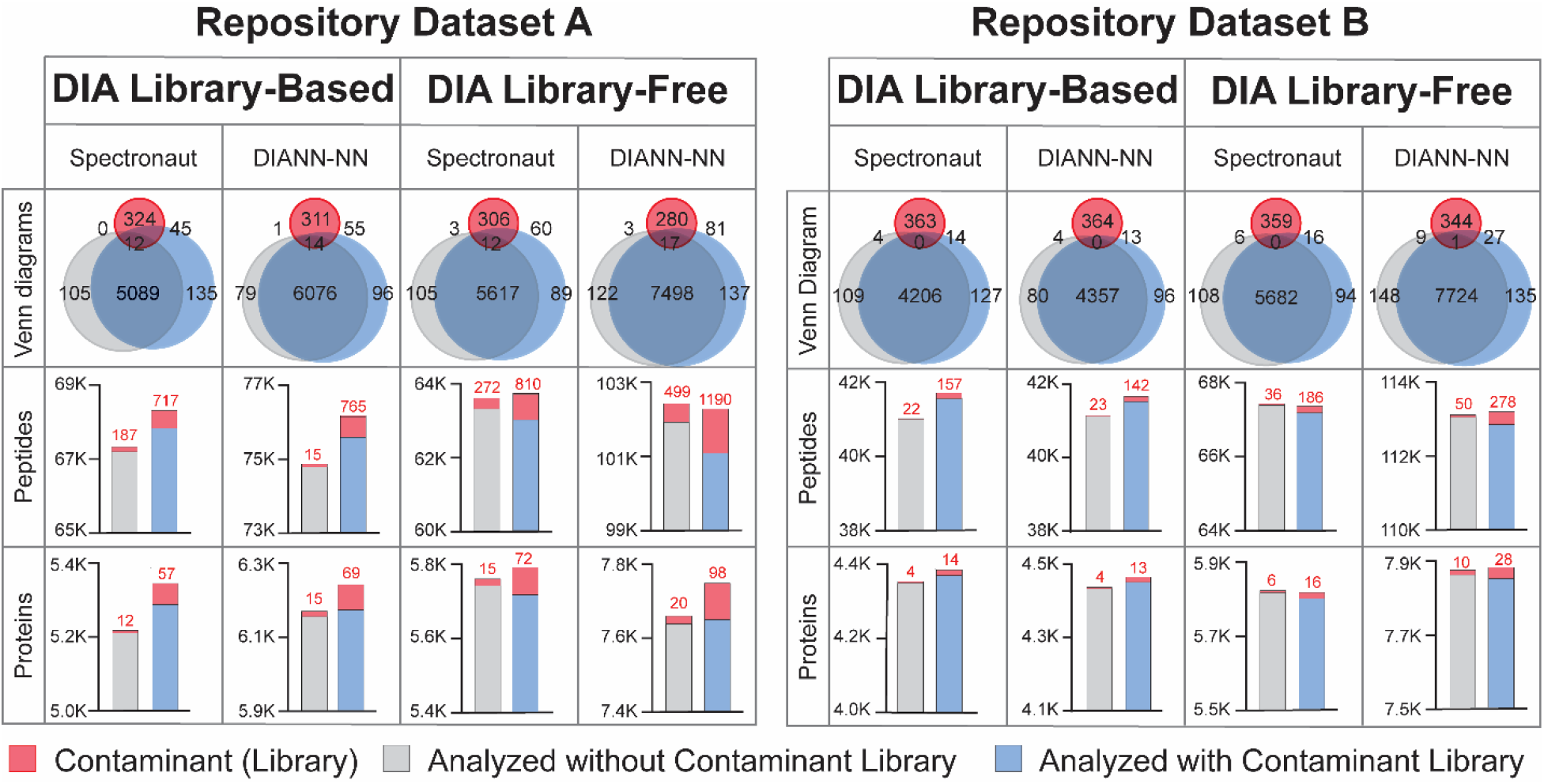
Repository DIA proteomic datasets reanalyzed with and without the contaminant protein libraries. Various DIA data analysis platforms were used to reanalyze two DIA datasets (human cells: PXD022589; mouse cortex: PXD005573) with (blue) and without contaminant libraries (grey). Contaminant FASTA library and identified contaminant proteins were marked in red in the Venn diagrams and bar graphs.

Since our contaminant libraries can improve protein/peptide identifications, we further assessed protein quantification with and without contaminant libraries. Coefficient of variation (CV) of all quantified proteins from HEK cells (**Figure 6A**) and mouse brain samples (**Figure 6B**) were calculated after removing the contaminant proteins. No significant differences were observed with and without including contaminant libraries. DIA-NN resulted in more protein identifications, but higher CVs compared to Spectronaut platform. Library-based methods provided less variation and better reproducibility compared to library-free methods, consistent with other reported studies.^23,45^ Protein intensities were not exactly the same when the data was analyzed with or without contaminant libraries, but they did correlate very well with spearman’s correlation close to 1 (**Figure 6C and 6D**). To rule out the possibilities that differences in protein identification and quantification may be caused by additional entries in the libraries, we performed a control analysis where 381 random proteins were removed from the human FASTA library (**Supplemental Figure S4**). Including contaminant library always outperformed the method without contaminant library. No major differences in quantification were observed in this control analysis, demonstrating that including additional contaminant libraries do not influence protein quantification.

**Figure 6:**
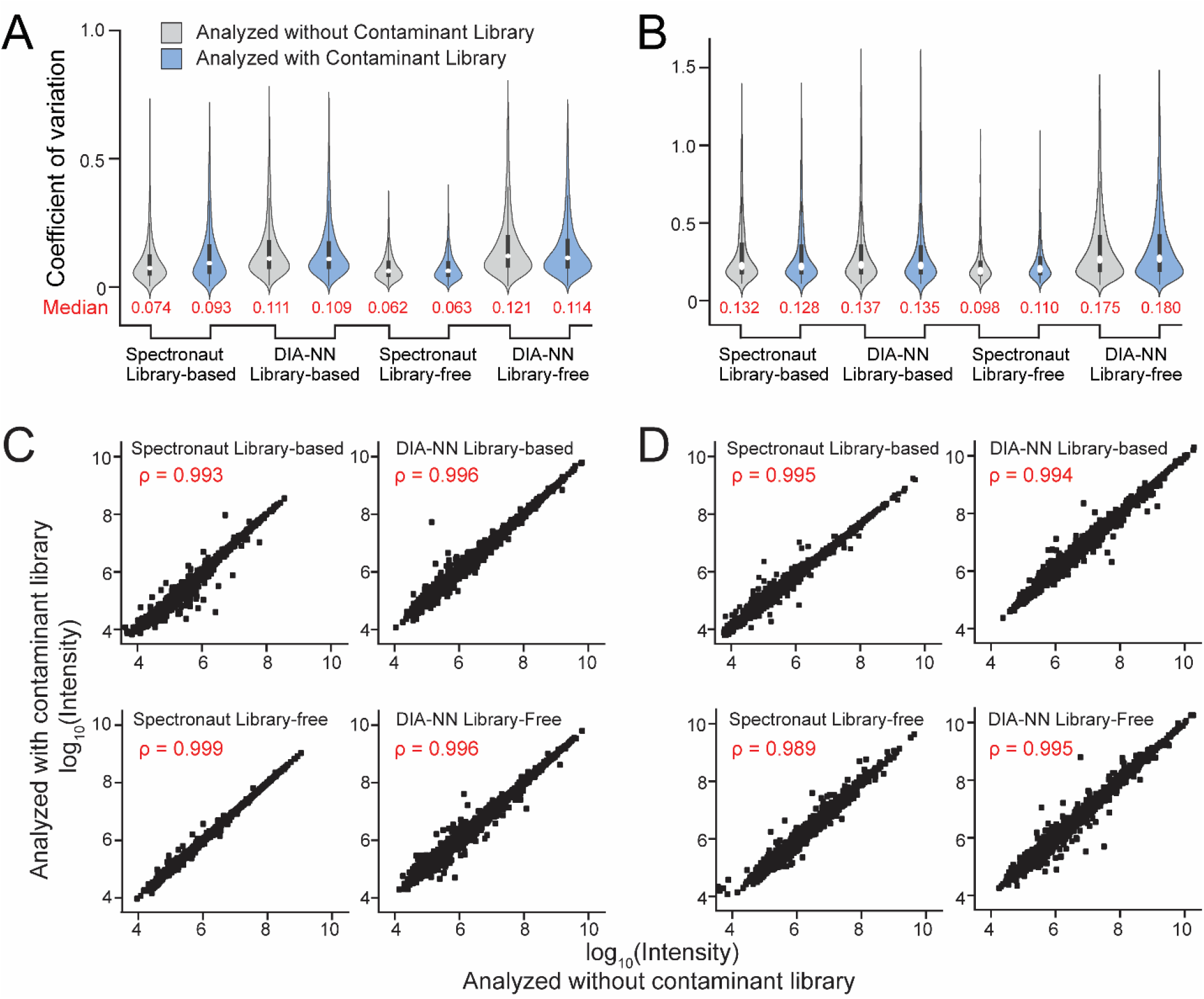
Protein Quantification is not Influenced by the Protein Contaminant Libraries. Violin boxplots showing the coefficient of variance for protein quantification with (blue) and without (grey) the contaminant library in HEK cells (A) and mouse brain tissue (B) DIA proteomics datasets. Spearman’s correlations of protein intensities were calculated with and without the contaminant libraries in HEK cells (C) and mouse brain tissue (D).

## CONCLUSIONS

To sum up, we highly recommend using our contaminant libraries for both DDA and DIA proteomics data analysis. This study filled a critical gap in bottom-up proteomics by establishing and evaluating contaminant protein libraries to reduce false discoveries and improve identifications in both DDA and DIA proteomics. Although the software used here (Spectronaut, DIA-NN, MaxQuant, Proteome Discoverer) are not an exhaustive list of all available data analysis platforms, we believe that our contaminant libraries can be universally applied to all bottom-up DIA and DDA proteomics software. In fact, we provided step-by-step tutorial on how to best incorporate our contaminant FASTA and Spectral libraries for many other software platforms such as Skyline^22^, MaxDIA^24^, and PECAN^27^ (**Supplemental Tutorial**). Recognizing the different nature of samples used in various proteomics experiments, our ongoing efforts will continue updating and enriching our contaminant libraries to include sample type-specific contaminant libraries on our website (https://github.com/HaoGroup-ProtContLib). Current available FASTA libraries include the universal contaminant FASTA evaluated in this study, as well as new FASTA libraries specifically for cell culture, mouse tissue, and rat tissue. These freely accessible contaminant FASTA and spectral libraries can be valuable resources for proteomics researchers and facilitate the standardization of proteomic data analysis across different laboratories.

## SUPPORTING INFORMATION

**Supplemental FASTA.** Contaminant protein FASTA with Cont_ prefix.

**Supplemental Tutorial.** Tutorial for using contaminant libraries for DDA and DIA data analysis in various proteomics software.

**Figure S1**: Additional examples of misidentified and coeluting contaminant peptides when not using contaminant library in DIA data analysis.

**Figure S2**: Evaluation of different methods to build contaminant spectral libraries.

**Figure S3:** Influence of protein contaminant libraries on identification and quantification in a controlled analysis to randomly remove 381 proteins from the human FASTA.

**Table S1**: Contaminant protein information in the FASTA library and sources of contaminations.

**Table S2**: List of the established FASTA and Pulsar spectral libraries.

**Table S3**: Summary of the protein contaminants identified in contaminant-only samples.

**Table S4**: List of the contaminant peptides and fragments in Pulsar spectral libraries.

## AUTHOR CONTRIBUTIONS

A.M.F. and L.H. designed the study and wrote the manuscript with inputs and revisions from all coauthors. A.M.F, J.N., and A.M. conducted the experiments. A.M.F and J.N., performed data analysis. All authors have read and agreed to the published version of this manuscript.

## ACKNOWLEDGEMENTS

This study is partly supported by the NIH grant (R01NS121608). L.H acknowledges the ORAU Ralph E. Powe Junior Faculty Enhancement Award. A.M.F acknowledges the ARCS-Metro Washington Chapter Scholarship and the Bourbon F. Scribner Endowment Fellowship. We thank the Vertes lab and Lu Lab at GW for the access to the SpeedVac equipment and mouse brain samples. We thank Tejas Gandhi and Oliver Bernhardt from the Biognosis Spectronaut team for providing valuable feedback on DIA data analysis.

---

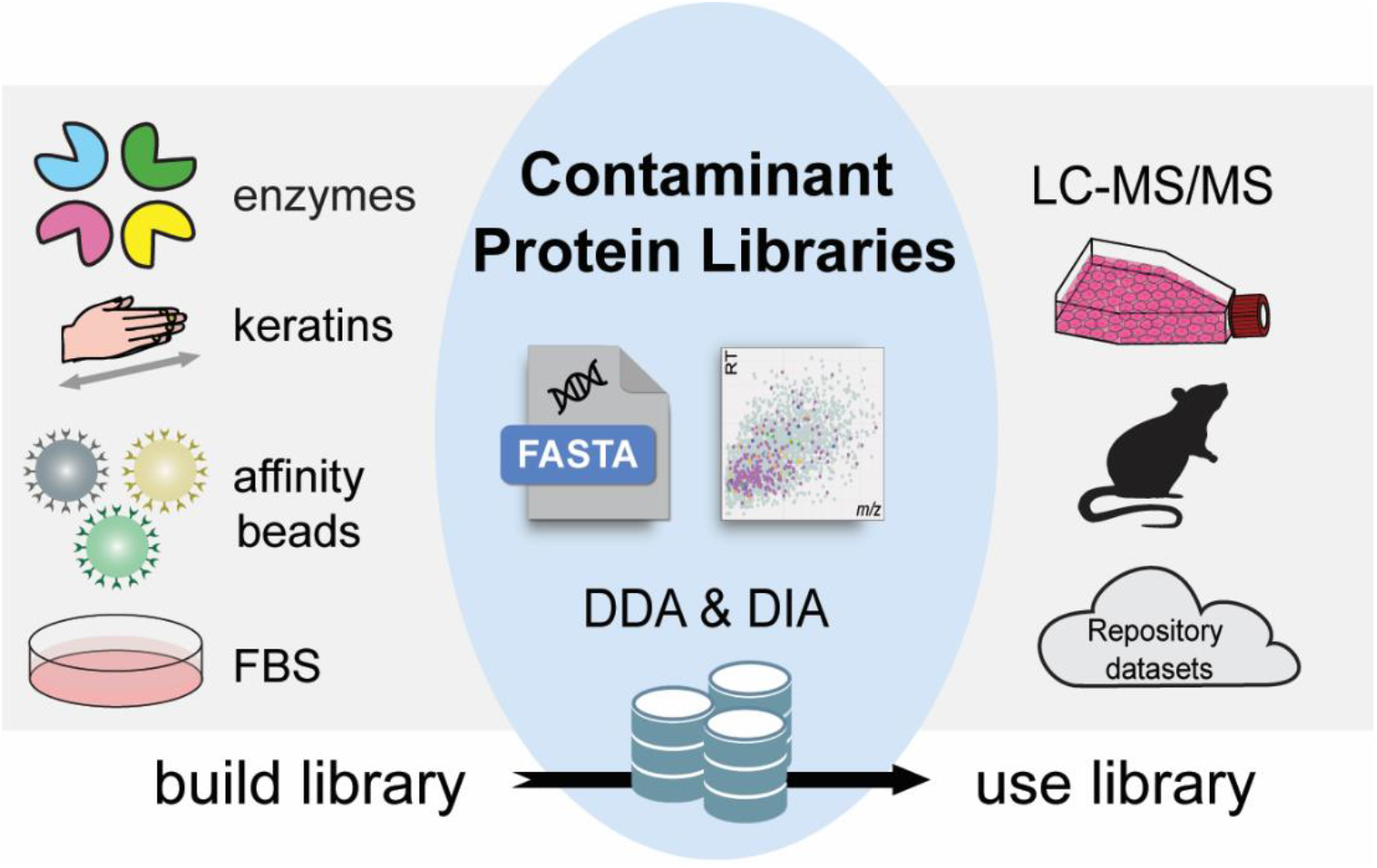

**Table of Contents (TOC) graphic**

(All images used in the TOC graphic are original, in compliance with copyright guidelines.)

