## Supporting Information for "Protein Contaminants Matter: Building Universal Protein Contaminant Libraries for DDA and DIA Proteomics"

Ling Hao

Assistant Professor

Department of Chemistry

George Washington University

### TABLE OF CONTENTS


- **Figure S1.** Additional examples of misidentified and coeluting contaminant peptides when not using contaminant library in DIA data analysis.
- **Figure S2**. Evaluation of different methods to build contaminant spectral libraries.
- **Figure S3**: Influence of protein contaminant libraries on identification and quantification in a controlled analysis to randomly remove 381 proteins from the human FASTA.
- **Table S1**: Contaminant protein information in the FASTA library and sources of contaminations.
- **Table S2**: List of the established FASTA and Pulsar spectral libraries.
- **Table S3**: Summary of the protein contaminants identified in contaminant-only samples.
- **Table S4**: List of the contaminant peptides and fragments in Pulsar spectral libraries.
- **Supplemental FASTA.** Contaminant protein FASTA with Cont_ prefix.
- **Supplemental Tutorial.** Tutorial for using contaminant libraries for DDA and DIA data analysis in various proteomics software.

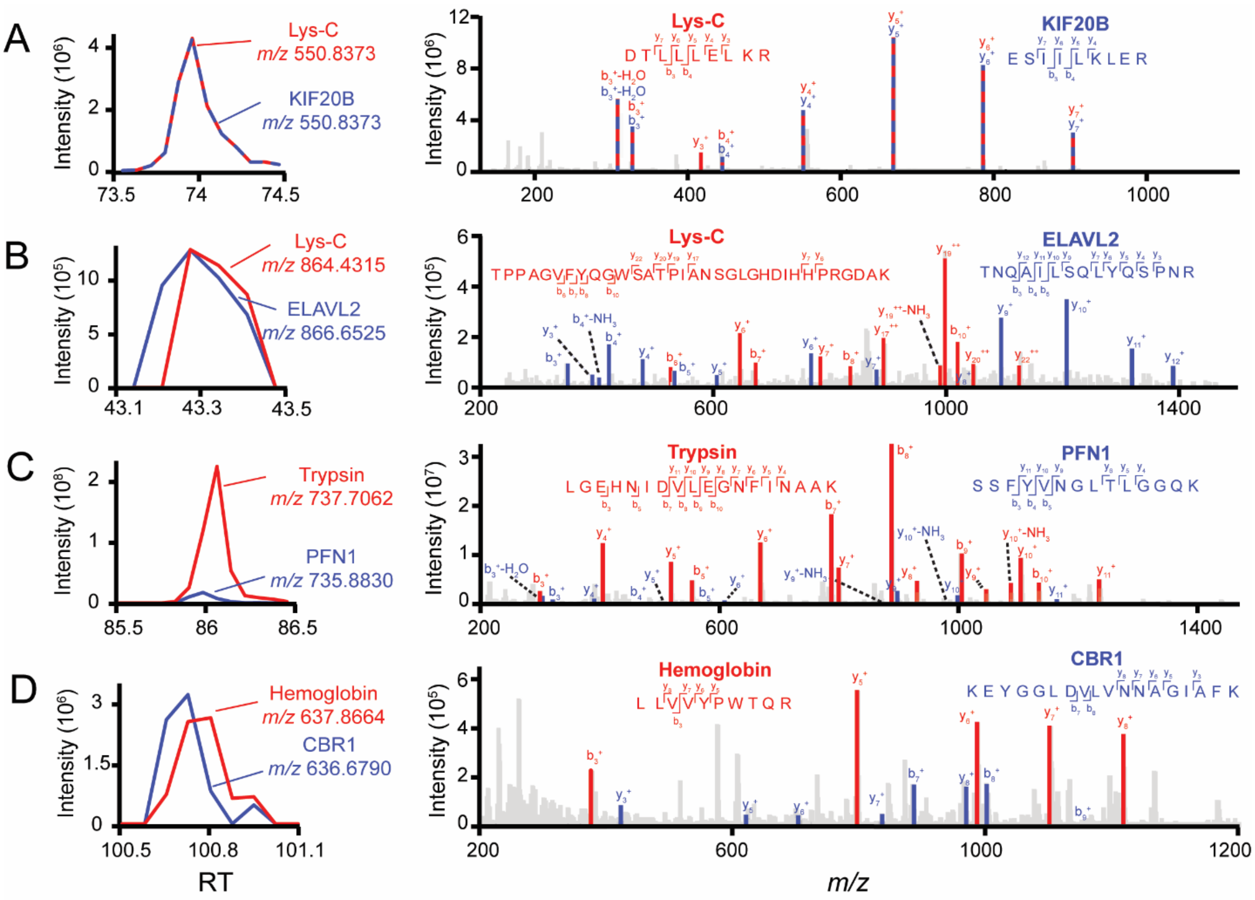


**Supplemental Figure S1: Additional Examples of Misidentified and Coeluting Contaminant Peptides when Contaminant Libraries are not Included for Data Analysis.** Example contaminant peptide chromatograms and MS/MS fragments were shown in red, real peptides of interest were shown in blue. (A) Lys-C (enzyme contaminant) was misidentified as KIF20B. (B) Lys-C was coeluted and co-fragmented with ELAVL2. (C) Trypsin was coeluted and co-fragmented with PFN1. (D) Hemoglobin was coeluted and co-fragmented with CBR1.

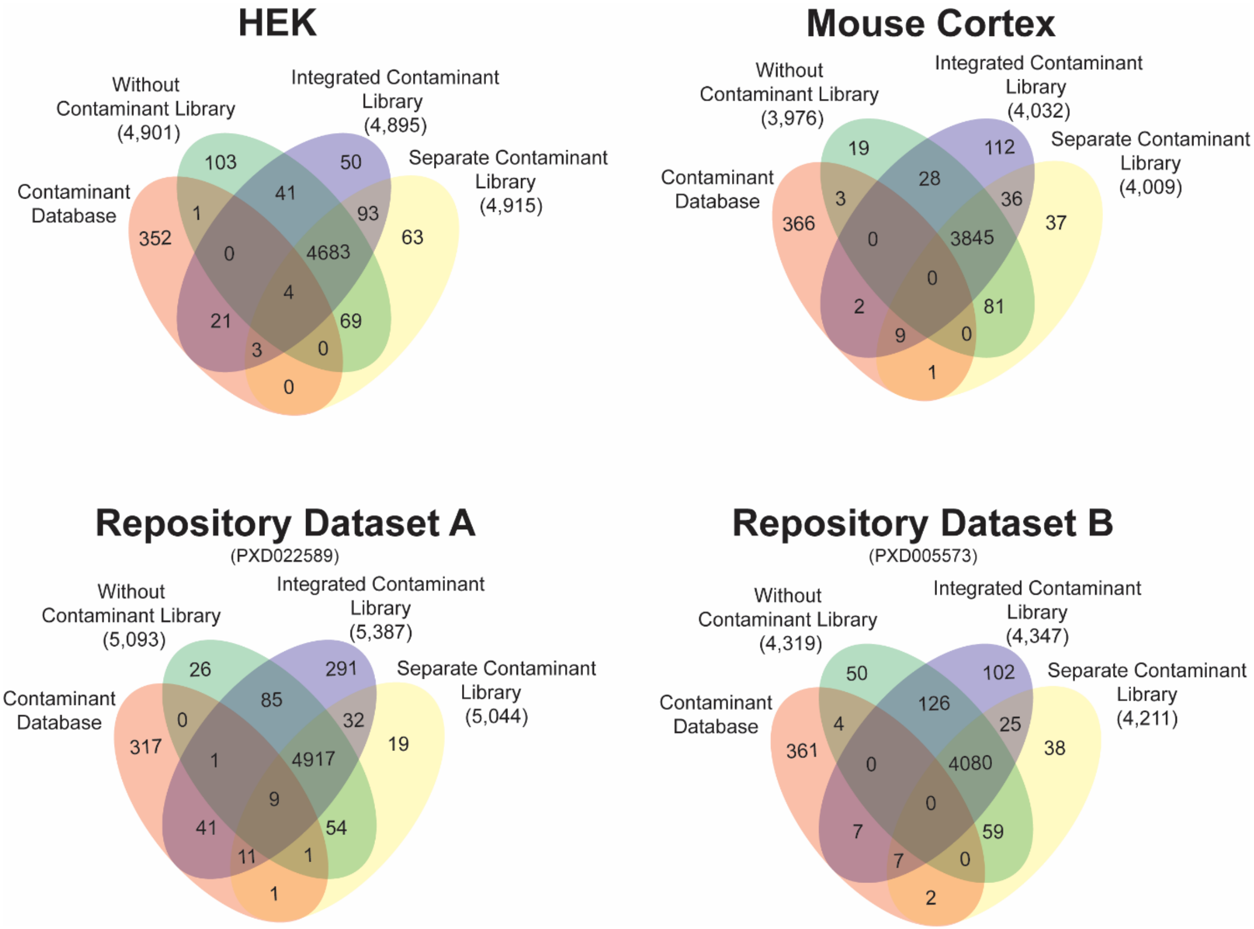


**Supplemental Figure S2**: **Evaluation of different methods to build contaminant spectral libraries.** Protein contaminant raw data files can be used to combine with the custom data to create one integrated library (purple) or included as an additional spectral library in the Spectronaut software (yellow), in comparison to the results without using contaminant library (green).

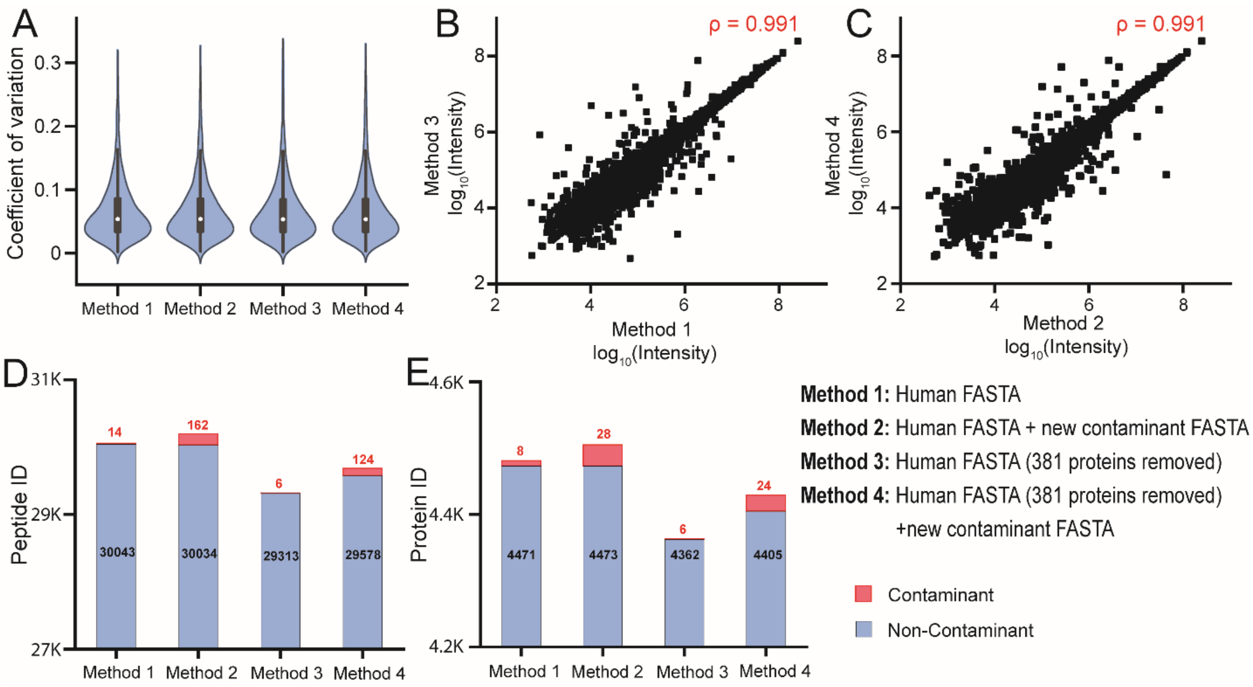


**Supplemental Figure S3: Influence of protein contaminant libraries on identification and quantification in a controlled analysis to randomly remove 381 proteins from the human FASTA.** The HEK dataset was re-analyzed using the modified Human FASTA with 381 proteins removed (Method 3), and the modified Human FASTA + new contaminant FASTA (Method 4). (A) The coefficient of variation is not affected by the addition of the contaminant FASTA. Spearman's correlations of peptide intensities were calculated for using different FASTA files (B, C). Comparison of DIA protein (D) and peptide (E) identifications from each method.
